# Tracking White Matter Changes After Stereotaxic Radiosurgery in Miniature Pigs Using structural MRI, DTI, and FDG-PET

**DOI:** 10.1101/2025.05.04.652148

**Authors:** Ke-Hsin Chen, Chun-I Yeh, Furen Xiao, Mei-Fang Cheng, Ruoh-Fang Yen, Yu-Ten Ju, Yilin Chen, John R. Adler, Daniel Barbosa, Michael Bret Schneider

## Abstract

Stereotaxic radiosurgery (SRS) non-invasively and precisely ablates brain tumors or glioma located in the location where is surgically inaccessible with the aid of three-dimensional coordination system. This technique can also treat functional or psychiatric disorders, yet its dosimetry and curative mechanism remain to be elucidated. In this study, a miniature pig model was utilized to verify the effect of stereotaxic radiosurgery on a white matter tract with various doses delivered by CyberKnife. As porcine brain bears high resemblance to that of human in size and structure, the potential irradiation-induced change in structures and metabolism was monitored by conventional tools in clinical, including anatomical magnetic resonance image (MRI), diffusion tensor image (DTI) and 18F-Fluoro-D-Glucose positron emission tomography (FDG-PET). The right internal capsule was selected as the surgical target, and a one-year longitudinal study was conducted with whole brain images obtained once per three months. The results indicate that a dose equal to or higher than 60 Gy led to a late-onset radionecrosis which took a period close to 180 d to develop edema and breakage in the blood-brain barrier. In the meanwhile, DTI indices and differential tractography further illustrate a dose- and distance-dependent white matter injury along the tract of internal capsule. In contrast, doses of 40 Gy and below did not result in any discernible harm to the brain structure, but a sustained local inhibition in brain metabolism was observed in some pigs. The modulatory effect of low dose radiation awaits a comprehensive assessment throughout the whole brain in combination with behavioral or cognitive tasks built on pigs. This study showing the dose- and time-dependent changes will improve the understanding of the SRS dosimetry on the white matter and help investigators to decide an optimal window for brain imaging or behavioral assessment to take place on patients.

## Introduction

Stereotaxic radiosurgery (SRS) applies ionizing radiation converging onto the target with the aid of three-dimensional coordination system, thus it can precisely ablate brain tumors or glioma in the position like regions consisting of elaborate structures or deep in the brain where is surgically inaccessible (Leksell, 1951, 1971). The cumulative high dose in the target can produce radiation damage, while the adjacent healthy tissues receiving a limited dose were spared. The radiation-induced ablation often takes more than 6 months to be observed, and is hypothesized to result from vascular damage or loss of glial cells (Greene-Schloesser et al., 2012; Tofilon & Fike, 2000). Therefore, a follow-up assessment of treatment response either in structural change or cognitive function is necessary to distinguish whether the patient is an SRS responder or not (Chen et al., 2017; Putz et al., 2020; Tohyama et al., 2018). As SRS is a non-invasive neurosurgical technique, multiple site could be treated or a site could be treated repeatedly with a fractionized dose prescribed according to the volume and type of the target. The duration for patients to be hospitalized can be significantly reduced as no incision is needed, therefore SRS becomes a popular technique for neurosurgery.

SRS can also treat functional or psychiatric disorders (Leveque et al., 2013), yet its dosimetry and curative mechanism remain to be elucidated. For example, patients with essential tremor (Tuleasca et al., 2021) or obsessive compulsive disorder (Miguel et al., 2019) were prescribed with a high dose eliciting necrosis (130-180 Gy at maximum) to disturb signal transmission in a specific neural pathway. A high dose ranging 60-90 Gy at maximum was also prescribed to the patients with trigeminal neuralgia (Tuleasca et al., 2019). As for the patients with drug-resistant refractory epilepsy, a moderate dose (20-25 Gy at the marginal isodose) would be applied for seizure control (McGonigal et al., 2014; McGonigal et al., 2017; Quigg et al., 2012). The curative mechanism of SRS in each disease still needs to further exploration, as radiation may evoke independent biological pathways to induce inhibitory and excitatory processes simultaneously in the target area (Di Veroli et al., 2015). For example, the cocade theory proposed by Regis et al. (2010) states that the cumulative high dose in the core region brings radiation necrosis while the relatively lower dose in the surrounding area exerts a neuromodulatory effect by increasing the inflammatory compounds without significant cell death. Furthermore, treated patients often report symptom relief earlier than the radiation-induced cellular or brain activity alteration which requires months to occur (Regis et al., 1999). Therefore, a longitudinal investigation on brain structure, neuronal activity, and behavioral assessment in combination of various radiation dose applying onto a specific target is necessary to delineate the SRS dose-response curve, and these results will help to unveil the mechanism of how SRS cures functional disorders.

The pig (*Sus scrofa*) is a large animal model with advantages in neuroscience research, especially in translational study. First, the structure, physiology and size of pig brain is similar to human brain (Lind et al., 2007; Sauleau et al., 2009). The porcine brain is gyrocephalic, which means cortices folding into gyri separating by sulci as circumvolutions (Bjarkam et al., 2017), whereas the rodent brain is smooth-surfaced.

The brain size of pig is bigger than that of rodent, and its scale is about one seventh of a human brain (180 vs 1,300 g) (Sauleau et al., 2009). As a matter of fact, the conventional tools in clinical, like neuroimaging techniques, radiosurgery, deep brain stimulation and so on, can be readily applied onto the porcine brain, and the target site can be pinpointed to a specific gyrus or a subcortical structure (Chang et al., 2021; Knight et al., 2013; Yeh et al., 2021; Zaer et al., 2020). Second, the porcine brain development bears high resemblance to that of human brain, thus pig model can serve as a good preclinical model for neurodevelopment (Conrad et al., 2012), neurodegenerative disease (Hoffe & Holahan, 2019) and aging (Schachtschneider et al., 2021). As pigs grow fast to attain sexual maturity at 6 months and have a life span of 12-15 years, a close investigation in pathological development can be conducted longitudinally within one cohort or in a cross-sectional fashion. The miniature pigs bred for biomedical purpose have advantages to be applied in the longitudinal study for their smaller body size and weight as compared with the agricultural breeds (Lind et al., 2007). Finally, the anatomical structures of pig brain have been described histologically (Bjarkam et al., 2017; Felix et al., 1999; Orlowski et al., 2019; Saikali et al., 2010), and brain templates built by magnetic resonance image (MRI) on various breeds of pigs were released in recent years (Chang et al., 2020; Conrad et al., 2014; Norris et al., 2021; Zhong et al., 2016). The connectome in structural or functional connectivity of pig brain has also been reported to have high degree of homology to that of human brain (Bech et al., 2020; Simchick et al., 2019; Zhong et al., 2016). These image results can promote the usage of pig brain especially in the field of neuroscience, and thus make miniature pigs become a popular model for human brain disorders.

Histology and brain imaging can help to decipher the effect of SRS in the central nervous system, as the former provides cellular information via microscopy and the latter shows brain signal change by specific image contrast. Postmortem tissues in concurrent with histological methods could reveal the SRS effect at the end point, yet this method was not able to depict the progress in structural or functional alteration unless a large sample size was recruited to be examined at specific time point (Tan et al., 2011). The non-invasive imaging techniques on the other hand can trace the progressive change on a subject during the follow-up period though a compromise in spatial resolution from micrometers to millimeters has to be reached. Nevertheless, the SRS effect can be evaluated via implementing proper image contrasts, such as monitoring brain metabolism by ^18^F-Fluoro-D-Glucose positron emission tomography (FDG-PET) (Yeh et al., 2021), locating edema and hemorrhage by MRI (Benczik et al., 2002; Kunimatsu et al., 2015) and so on. When radiosurgery was applied onto neural fibers, diffusion tensor image (DTI) which infers plausible fiber orientation by a tensor model depicting the anisotropy of water diffusion can provide quantitative indices to measure the white matter microstructure (Yang et al., 2021; Zhang et al., 2012), and these indices can further be correlated with the medical outcome to examine the clinical effectiveness. For example, the decrement in fractional anisotropy (FA), an index reflecting the integrity of white matter, detected 6 months after SRS could prognosticate long-term pain relief on patients with trigeminal neuralgia (Tohyama et al., 2018). Furthermore, tract segments with FA decreasing mapped by differential tractography from repeat DTI scans can serve as a biomarker indicating white matter injury (Yeh et al., 2019). This novel tractography technique has not yet been used in the SRS research but has a promising potential to provide diagnostic and prognostic evaluation at individual level.

The purpose of this study aimed to use porcine brain as a translational model to evaluate the effect of SRS on the white matter tract with various radiation doses, from a necrotic high level to a minimal one. The internal capsule (IC) was selected as the target for its stalk (about 4∼5 mm) is larger than that of the corpus callosum (less than 1.5 mm) in the coronal section of the pig brain, and a collimator of 5 mm can produce radiation converging onto this tract. A one-year longitudinal study including a pre-surgery baseline and follow-up brain scans once per three months was adopted to reveal alterations in brain structures and metabolism by MRI and FDG-PET, respectively. In the meanwhile, white matter microstructure and tract segments showing radiation-induced injury were investigated by DTI in combination with differential tractography. The results of this study showing the dose- and time-dependent changes will improve the understanding of the SRS dosimetry on the white matter and help investigators to decide an optimal window for brain imaging or behavioral assessment to take place on patients receiving SRS.

## Methods and Materials

### animals

Lee-Sung (LS) miniature pigs with the average of 5.6 ± 0.9 months old and 19.8 ± 2.1 kg were recruited in this study. The white-coated LS pig was a selective breed by mating Lanyu sows, indigenous black-coated miniature pigs in Taiwan, with Landrace boars (75% Lanyu and 25% Landrace) in Department of animal science and technology, National Taiwan university. Eight LS pigs (1 female, 7 male) were randomly assigned to receive different SRS dose on two brain sites: the left primary motor cortex (L-M1, 10- 120 Gy) and the right internal capsule (R-IC, 2.5-100 Gy), and three animals (1 female, 2 male) were recruited to serve as non-irradiated control. All experimental procedures were conducted in accordance to the guidelines of the United States Department of Agriculture and were approved by the University Animal Welfare Committee in National Taiwan University.

### anesthesia and stereotaxic radiosurgery

The detail of experimental procedures can be referred in the work of Yeh et al. (2021) which had published part of the data, and a concise version would be recapitulated in the following. Animals were fasted for at least 12 hours to be anesthetized before the experimental procedures, including fiducial marker implantation, brain scans, as well as radiosurgery. The anesthesia was initiated by an intramuscular injection of Zoletil (4 mg/kg, Virbac Laboratories, Carros, France) and xylazine (2 mg/kg, Rompun, Bayer Corp., Whippany, NJ, USA) and then was maintained with 1-1.5 % isoflurane (Forane®, Arkema, Inc., Colombes, France) vaporized in the air through an orotracheal tube to control respiration rate at 0.2 Hz. The isoflurane was turned off at the end of experiment, and a single shot of tolazoline (2 mg/kg, Tolazine®, Akorn, Inc., Lake Forest, IL, USA) was given intravenously to curtail the anesthetization. Mechanical ventilation was used till the animal can maintain spontaneous breathing.

Vital signs including heart rate and blood oxygen level were closely monitored during the whole period of animal transportation and experiment, and blood sugar level was measured twice, before and after PET scan.

Three weeks before SRS, four titanium screws (Synthes Maxillofacial system, Synthes GmbH, Oberdorf, Switzerland) were implanted into the skull as fiducial markers. According to the baseline scan of MRI and positron emission tomography-computed tomography (PET-CT), the target definition and radiosurgical treatment were planned by the MultiPlan system version 2.2.0 (Accuray, Inc., Sunnyvale, CA, USA) with fiducial tracking model. On the day for radiosurgery, the anesthetized animal covering with a custom thermoplastic mask was placed in prone position on the table. An effective collimation diameter of 7.5 mm at the isocenter for L-M1 and 5 mm for R-IC was selected, and irradiation was prescribed to the 80% isodose line and the maximum dose (Dmax) as 100% by the CyberKnife system.

### image acquisition

Brain images obtained before SRS served as the baseline, and follow-up scans were obtained within a week (denoted as 7 d) and at 90, 180, 270, and 360 d after SRS. As for the non-irradiated control animals, brain images were acquired every 3 months (90, 180, 270, 360 d) after the baseline scan. MRI scans were acquired by the Achieva 3 T scanner (Philips, Eindhoven, The Netherlands) with an 8-channel body surface coil and the technique SENSE was implemented for parallel imaging (Pruessmann et al., 1999). A total duration about one hour was needed for obtaining anatomical images with various contrasts as well as DTI. The field of view (FOV) of all images was set as 200 x 200 x 90 mm to cover whole pig brain. A T2-weighted (T2W) image was obtained by using the turbo spin echo (TSE) sequence with repetition time (TR) = 3000 ms, echo time (TE) = 90 ms, flip angle (FA) = 90 degree, with a reconstruction matrix 480 x 480 and 90 slices. An image of fluid-attenuated inversion recovery (FLAIR) was used to detect edema in tissues with prolonged T2 relaxation time while suppressing cerebrospinal fluid (CSF) signal by the parameters of TR/TE = 9000/125 ms, FA = 90, inversion time = 2500 ms, NEX = 2, reconstructed matrix size = 480 x 480, slice thickness = 4 mm. T1-weighted images before and after a single dose of gadolinium (0.1 mmol/kg) were obtained (T1W and T1W-Gd, respectively) by using the fast field echo (FFE) sequence with TR/TE = 32/1.9 ms, FA = 30 degree, NEX = 1, matrix = 256 x 256, and slice thickness = 1 mm, while the T1W was only used for CT-MRI registration as its contrast not being able to detect subtle SRS-induced change. DTI was acquired by a spin-echo single-shot echo-planar imaging sequence with a b-value of 1000 s/mm^2^ along 32 directions plus one image without diffusion weighting (Bo), TR/TE = 2746/87 ms, NEX = 2, matrix = 128 x 128, and slice thickness = 2 mm.

PET-CT images were obtained by the Discovery PET-CT 710 scanner (GE Healthcare, Milwaukee, WI, USA), and the metabolic tracer ^18^F-Fluoro-D-Glucose (FDG) was administrated with 5-7 MBq/kg when the animal was maintained under 1% isoflurane in a quiet, dimly lit room. A period of 40 mins was allowed for the tracer to distribute in the pig’s body and then 30 mins were required for image acquisition. The FOV was set as 400 x 400 mm in the coronal view of pig brain, and a matrix size of 512 x 512 and 47 contiguous slices with 3.27 mm-thick were set for CT image, while the matrix size changing to 256 x 256 for PET.

### anatomical MRI and PET-CT analysis

The anatomical MRI and PET-CT images were pre-processed and analyzed by AFNI (Cox, 1996, 2012). All the MR DICOM files were first converted into NIfTI format by MRIcroGL/dcm2niix (Li et al., 2016), while the software 3D slicer (Fedorov et al., 2012) was used to calculate the PET standardized uptake values (SUV) and export files into NIfTI format. The image orientation of all images was well-checked via the marker put in the left side during image acquisition. The anatomical MR images with different contrast were first registered to the T2W image which had best resolution and image quality, and then were clipped to remove structures other than the brain and skull. Through rigid body transformation with normalized mutual information as the cost function (Maes et al., 1997), the CT image was registered to the T1W image of which the voxel intensity was inversed to resemble CT contrast. The PET SUV images were up-sampled to match the dimension of CT by trilinear interpolation and then transferred to the MRI space after visual inspection of CT-T1W registration results. The brain was retrieved by AFNI 3dSkullstrip in combination with manual modification to make a mask covering the whole brain excluding the olfactory bulbs.

Region of interests (ROI) analysis was adopted to reveal the signal alteration along the one-year follow-up period. According to the treatment plan in the baseline scan of MRI, a 5-mm-radius sphere was selected as ROI in the planned center at the right internal capsule (R-IC) to cover 80% isodose line, and a mirrored homologue was put in the non-treated left site (i.e., L-IC). These two IC ROIs were registered from the baseline to each of the latter scans via AFNI @animal_warper, a nonlinear image transformation method designed specifically for animal brain (Jung et al., 2021), in order to accommodate natural brain growth. The IC ROIs for non-irradiated animals were built with the same manner but the representative center was defined according to the cumulative map of irradiation overlaying on an average LS pig brain (depicted later). The signal ratio that serves as the index denoting SRS-induced MRI signal and PET SUV change was calculated by dividing the mean value of the R-IC from that of the L-IC, a reference ROI where is assumed to have negligible signal change with time. The derived ratio at each time point was further normalized by the pre-surgery one and then expressed in percentage change. With this normalization step the comparison in one dose across time or various doses at a specific time point is possible as it renders MRI original signals into scores in ratio scale and controls the basal activity in FDG PET. As the original IC ratio in the pre-surgery baseline approximates 1, a relative percentage change close to 0 % at a time point is interpreted as the signal intensity was comparable to the baseline, while a score higher or less than 0 % denotes the R-IC signal being enhanced or reduced, respectively.

### DTI analysis

DTI dataset were preprocessed with FSL (Jenkinson et al., 2012) to correct eddy current-induced distortion and head movement (Andersson & Sotiropoulos, 2016), and then diffusion tensors were fitted at each voxel to calculate scalar maps of fractional anisotropy (FA), mean diffusivity (MD), radial diffusivity (RD), and axial diffusivity (AD). Considering the pneumatic cavities within skull enlarged with pig age and no reversed phase-encoding dataset was obtained in the present data for susceptibility correction, the pre-surgery Bo image showing least distortion was first registered to the T2W image and then an inverted transformation matrix was obtained to warp IC ROIs into the DTI space. The IC ROIs were registered from baseline to latter time points in each pig nonlinearly between FA maps, and the value of diffusivity matrices was calculated after visual inspection of ROI location. The change in these quantitative matrices of the R-IC was expressed in percentage relative to the value of pre-surgery baseline to trace possible SRS-induced alteration at each follow-up time point, and the results of L-IC served as the negative control.

Tractography were conducted by DSI studio (http://dsi-studio.labsolver.org). In the conventional tractography, the fiber tracts passing through the region planned to receive SRS were delineated by setting the R-IC ROI as the seed region, the threshold for anisotropy and angular degree at 0.2 and 70, respectively, as well as regions of avoidance including the contralateral hemisphere and cerebellum, To map the segment of tracts showing axonal injury by differential tractography (Yeh et al., 2019), the image data of 7, 90, 180, 270, 360 d were spatially aligned to the pre-surgery baseline to calculate the difference in the anisotropic component of the spin distribution function (Yeh et al., 2010). After passing quality control test and having the goodness-of-fit between images evaluated by the software (R^2^ > 0.63), all warped images were visually inspected and two data were removed from analysis due to unsuccessful alignment (40 Gy at 270 d, 2.5 Gy at 360 d). The tracts with FA decrement were mapped by placing a total of 1,000,000 seeds in the whole brain excluding the cerebellum, and default value was used in the angular threshold, step size, and anisotropy threshold. The results obtained from 7d in the SRS-treated pig were used to estimate the false discovery rate (FDR) under the assumption that there should be no obvious structural change during this short course thereby any tracts showing FA decrement can be regarded as false-positive findings. To set a common criterion for differential tracking for all subjects, the pig receiving 100 Gy in the R-IC was chosen to test the FA change threshold (10 to 40 % with 5 % as an interval) and length threshold (5, 10, 15, 20, 25 mm) as its MRI data illustrate obvious tissue damage at 360 d but no change at 7 d (**Table S1**). A conservative FDR smaller than 0.01 was selected by setting FA decrement more than 30 % and tract length longer than 10 mm in the whole brain tracking, and the R-IC ROI were used to filter the tracts-with-difference passing through.

### An average brain of Lee Sung miniature pigs

An average brain of LS pigs (n = 11, denoted as LS11) was built to create representative IC ROIs for non-irradiated pigs and to illustrate the regions receiving SRS treatment in the coordination matched to the published atlases. A two-step registration-signal averaging procedure with a nonlinear transformation method was adopt to build the average T2W and DTI-FA images, by which improvement in signal-to-noise ratio and reduction in intra-subject variability were achieved (Holmes et al., 1998; Jung et al., 2021; Norris et al., 2021). The images from baseline scans were first registered to a subject selected arbitrarily to create an intermediate image by averaging the aligned results, and this mean image was used as the target for registration in the second iteration to create the final average brain. The ROIs of the planned SRS target were warped into the standard space to build a cumulative map illustrating the region subjected to irradiation across 8 pigs, and the voxel with peak value in this map was chosen as the representative center to create IC ROIs for control animals. The image coordination in the LS11 brain was the same as the atlas reported by Saikali et al. (2010) with the x, y, z axis corresponds to the mediolateral, rostrocaudal, dorsoventral axis, respectively. The AC-PC axis was in the midsagittal plane with the posterior commissure (PC), but not the anterior commissure (AC), serving as the zero point in the Cartesian coordinate system, with which is consistent with what used in the published pig brain template (Chang et al., 2020; Felix et al., 1999; Saikali et al., 2010; Zhong et al., 2016).

## Results

Figure 1 illustrates the region of the R-IC subjecting to SRS in a custom-made LS pig average brain with the PC as the zero point in the coordination system. The cumulative map overlaying on the T2W and FA images was made by registering the ROIs covering the planned SRS target with 80% iso-dose line to the standard space, and its number denotes how many times a representative voxel was chosen to be irradiated across 8 pigs. The treated area of all pigs centralized in the stalk of the R-IC, except for one target site (100 Gy) located more posteriorly and dorsally but this did not influence the explanation for the obtained results.

**Figure 1.**
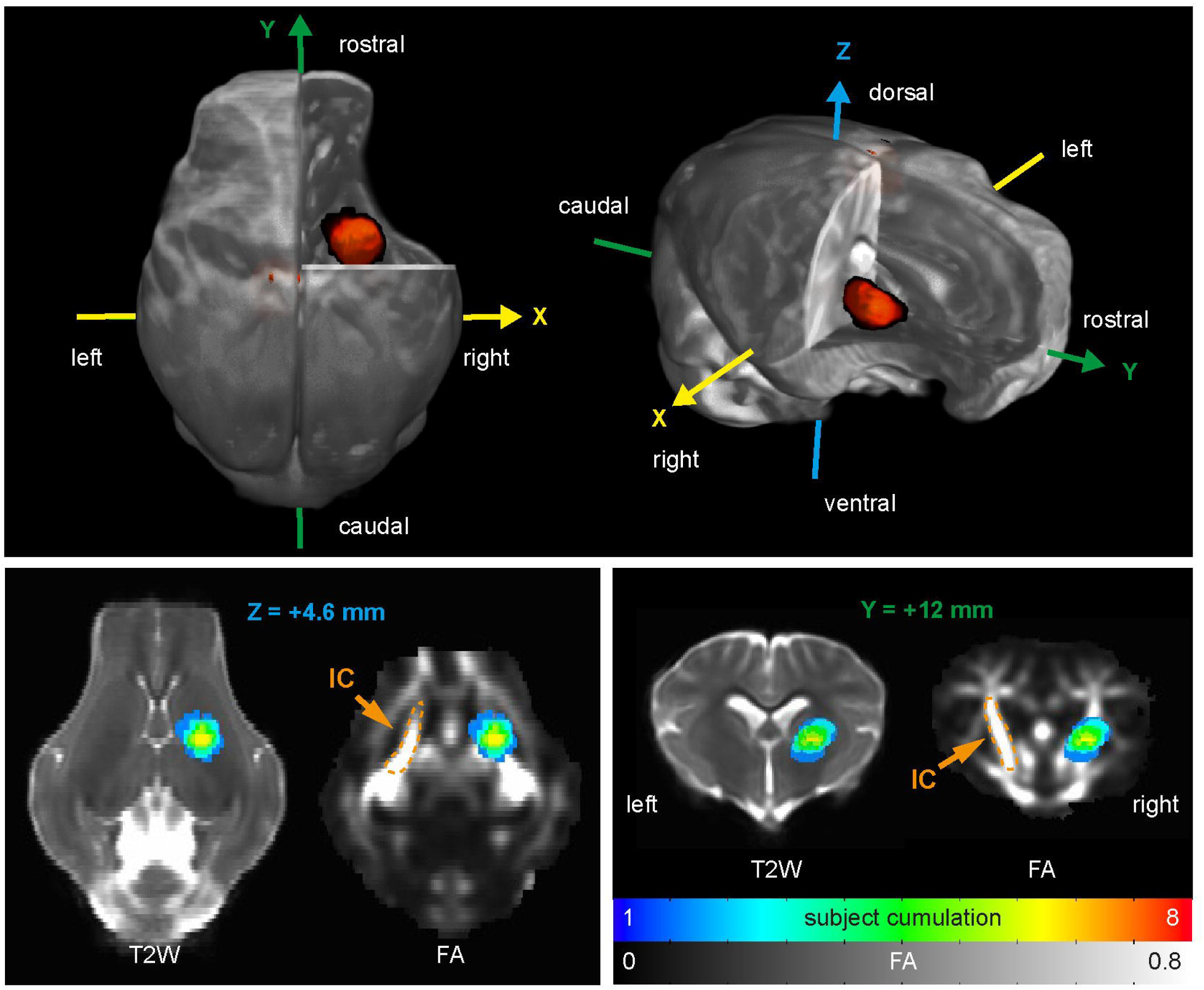
The stereotactic radiosurgery target site in the right internal capsule. (Top panel) The average Lee-Sung pig brain adopted the coordination system with the posterior commissure (PC) serving as the zero point, and the surgery target shown in red was located in the R-IC. (Bottom panel) The color of the cumulative map overlaying on horizontal and coronal slices of the T2W and DTI-FA denotes the number of subjects receiving radiosurgery in a representative voxel, and the peak value locates in the stalk of the R- IC. The slice position is illustrated in the distance (mm) relative to the zero point of the coordination system, and the L-IC in the DTI-FA image is highlighted for demonstration. IC for internal capsule.

Figure 2A shows that the SRS-induced R-IC signal alteration was found in the pigs receiving the maximum dose (Dmax) of 100 and 60 Gy and a period close to 180 d was needed for brain change to occur. In contrast, no obvious change was detected in the pigs receiving a dose of 40 Gy or below. The photo montage was made to emphasize the change in the surgical site by aligning follow-up images to the baseline scan through rigid-body transformation, by which only translates and rotates the brain images. The voxels with signal intensity over 95^th^ percentile rank in the FLAIR and over 80^th^ in the T1W-Gd, chosen arbitrarily, were colorized and overlapped onto the T2W for the ease to trace the pathological progress. A perifocal edema and radiation necrosis were observed at 180 d in pigs receiving 100 and 60 Gy, with blue color denoting increased fluid content around the surgery center and red color denoting an increased gadolinium uptake via broken blood-brain barrier (BBB). These pathological changes gradually recovered at 270 d and 360 d, but hematomas were noted with hypointense in T2W and FLAIR of the pig receiving 100 Gy. The dose of 100 Gy further caused brain structure deformation and right ventricle enlargement in the long run. As for FDG-PET images in Figure 2B, a slight SUV enhancement found in R-IC than L-IC at 180d for 60 and 100 Gy did not reflect brain activity increment per se but rather a local blood leakage as indicated by T1W-Gd, whereas the R-IC SUV in lower dose bear no relationship to BBB damage. It is worth noting that the basal brain activity might be various among pigs under the condition that the L-M1 and R-IC were manipulated simultaneously for parsimony in subject number. The R-IC to L-IC signal ratio relative to the pre-surgery baseline level was implemented to enable comparison across time and subject in the following ROI analysis.

**Figure 2.**
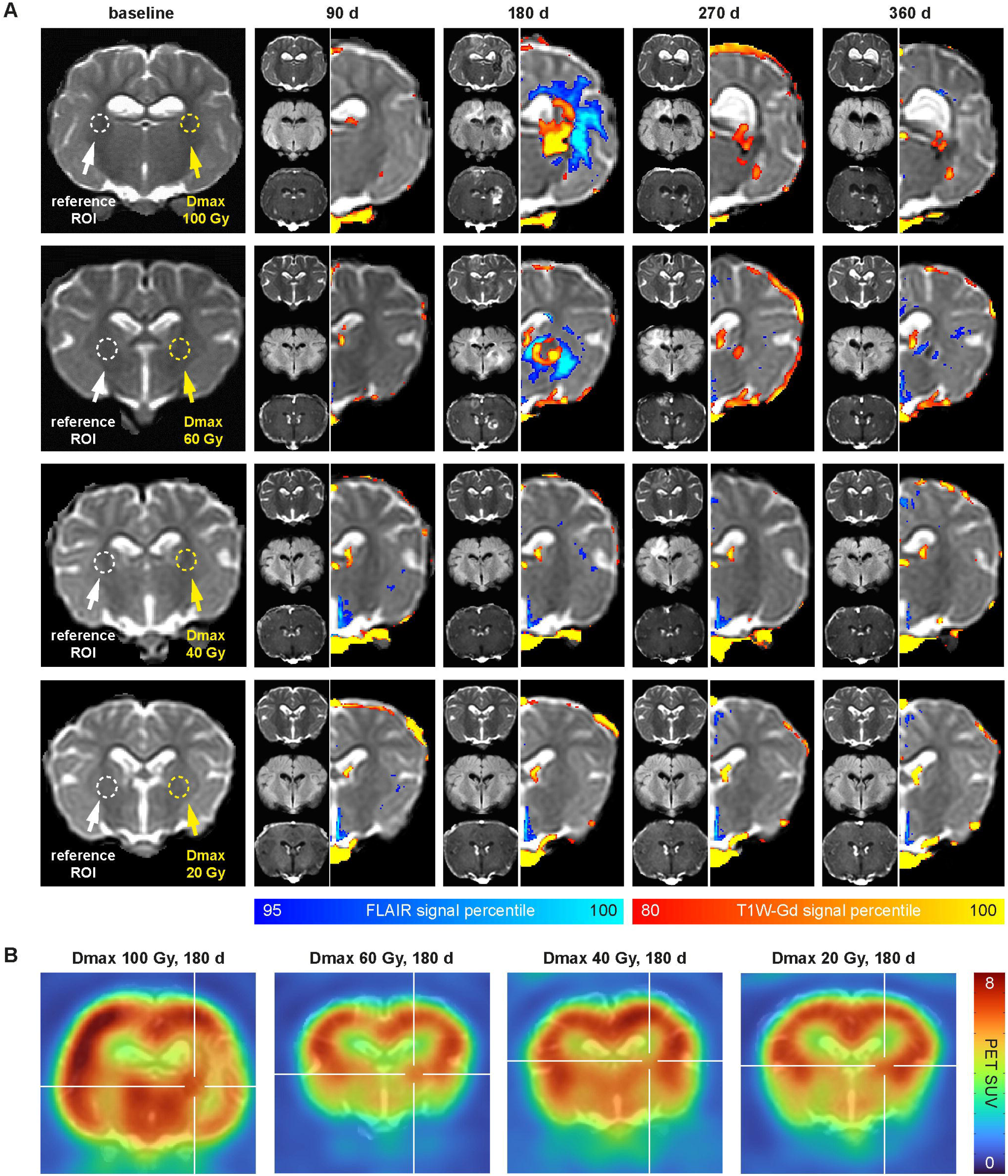
The radiation-induced brain change was found 180 days after SRS when a dose of 100 or 60 Gy was delivered. (A) The pre-surgery baseline image illustrates the surgical site on the R-IC with the maximum dose (Dmax) and the reference ROI on the L-IC. For images obtained in the follow-up period, the T2W, FLAIR and T1W-Gd from top to bottom were shown in the left side, and voxels with signal intensity over 95th and 80th in the latter two, respectively, were colorized and overlapped onto the T2W image to emphasize the SRS-induced edema and necrosis in the right hemisphere. (B) The FDG PET images with color indicating SUV were overlapped on the T2W, and the surgical site on the R-IC was high-lighted by a white cross. A slight SUV enhancement was found in the pigs receiving SRS dose 100 and 60 Gy at 180 d when being compared with their L-IC, but it did not reflect brain activity increment per se but rather a blood leakage from the damaged BBB as indicated by T1W-Gd in Figure 2A.

The ROI analyses in Figure 3 recapitulate the SRS-induced brain alteration by the IC signal ratio relative percentage change, and the data from three non-irradiated pigs were averaged to become a representative subject for negative control (0 Gy, black line). The result first confirms that no significant MRI signal alteration was observed in the Dmax dose of 40 Gy or below in all follow-up scans. Secondly, a period close to 180 d was necessary for SRS effect on brain structure to occur, as a magnificent signal elevation could be detected from T2W, FLAIR, and T1W-Gd images especially in 60 Gy. As for the necrotic dose 100 Gy, the signal value was comparable to normal tissue in T2W and FLAIR at 180 d owing to the mixture of edema and necrotic lesion within the R- IC ROI and it became lower at 270 and 360 d due to the hypointense foci indicating hematoma. On the contrary, signal of Gd-enhancement clearly indicated BBB impairment at 180 d while a gradual recovery was found in later time points by showing a mild signal elevation in both 100 and 60 Gy. Finally, low dose radiation may have a potential in neuromodulation as a small yet stable inhibitory effect was found in some cases by FDG-PET. Being compared with the negative control who had an unchanged IC ratio across time, the ratio of treated subjects did have a small change of 10 %, a level was also reported in the L-M1 (Yeh et al., 2021). Although the dose of 20, 10 and 5 Gy had no clear impact, the dose 40, 30, and 2.5 Gy exerted a long-term inhibition on focal metabolism lasting a half year from 180 d. Further, a negative correlation was found between the SRS dose and IC ratio percentage change at 360 d (*r*(5) = -.768, *p* = .04), where 100 and 60 Gy were excluded from analysis for their necrotic effect. Further investigation is needed to ensure the SRS modulatory effect on the white matter by recruiting more subjects in the future.

**Figure 3.**
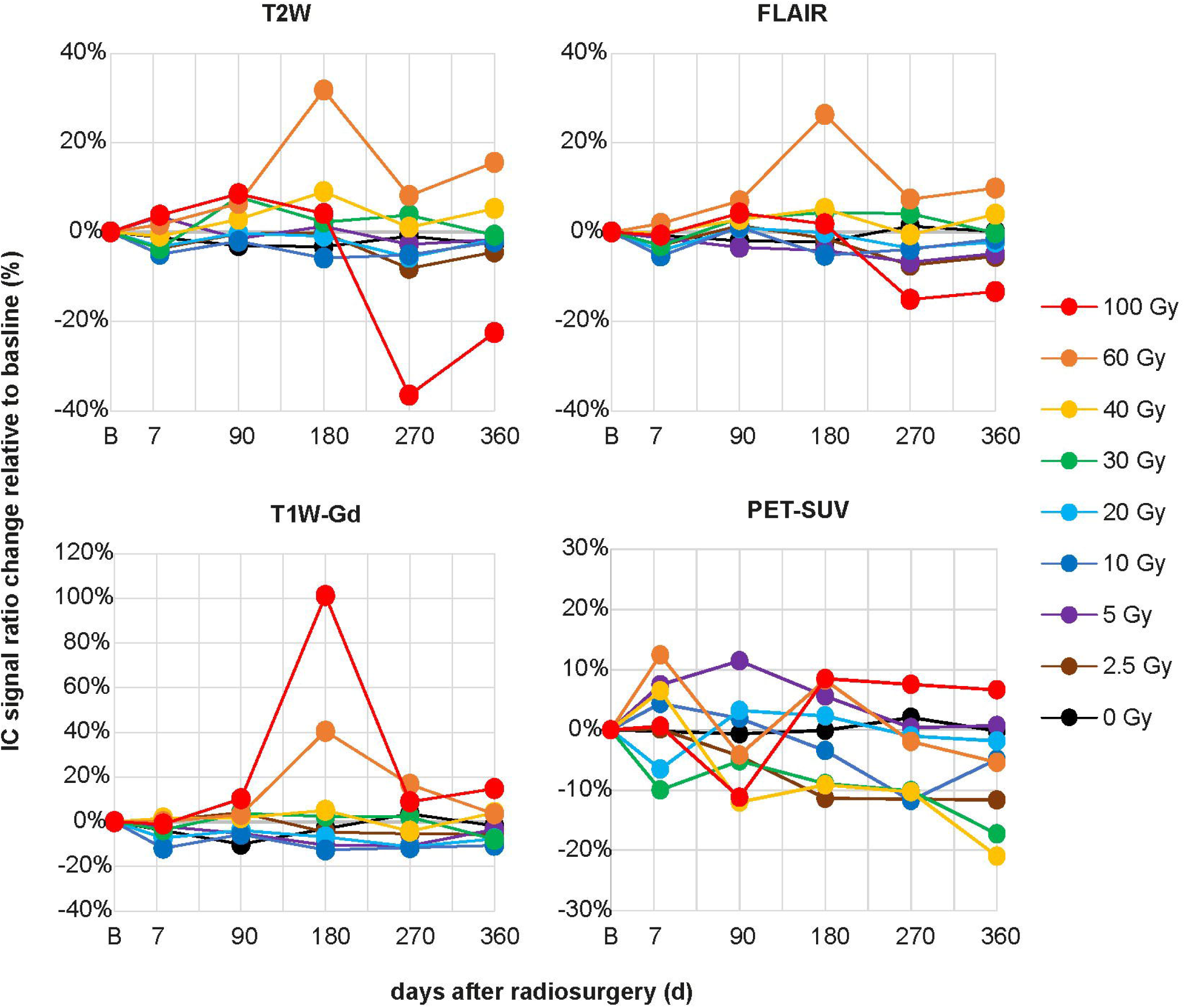
ROI analyses on T2W, FLAIR, T1W-Gd, and FDG-PET. ROI analyses from MRI show a period close 180 d was necessary for pathological change such as edema and radiation necrosis to occur when SRS dose of 100 or 60 Gy was applied. On the contrary, no obvious signal change was detected in the pigs receiving a dose of 40 Gy or below. The result from FDG-PET indicates the dose of 40, 30, and 2.5 Gy exerted a long-term inhibitory effect from 180 d with which a potential radiosurgery-induced neuromodulary effect was observed. The data from control animals (n = 3) were averaged to become a representative subject (0 Gy, black line). A normalized IC signal ratio relative change higher or lower than 0% indicates the R-IC signal being respectively elevated or decreased in comparison with the pre-surgery baseline denoted by B in the x axis.

The ROI analysis of DTI quantitative indices showed a congruent result that SRS-induced white matter damage occurred in the dose equal to or higher than 60 Gy and this effect was detected at 180 d after surgery. Figure 4A illustrates all DTI indices of R- IC percentage change relative to pre-surgery baseline, and the original value was shown in **Figure S1** with L-IC serving as negative control. An FA decrement more than 40% in 60 and 100 Gy-treated R-IC was found from 180 d persisting to 360 d, indicating a high radiation dose caused a white matter damage as the degree of anisotropy in water diffusion restricted by myelin was impaired. MD reflects mean water diffusivity regardless of the direction, while RD increment and AD decrement are sensitive to the status of demyelination and axonal degradation (Sun et al., 2006; Yano et al., 2018), respectively, by the extent of diffusivity perpendicular and parallel to the main axis of the tensor ellipsoid. In the case of Dmax 60 Gy, an AD decrement appeared 90 days earlier than the increment in RD and MD detected at 270 d, suggesting the injury of white matter came with axonal degradation first and then demyelination later. The slightly elevated AD at 360 d was uncoupled to axonal regeneration but rather related to a chronic injury showing mild edema around the surgery center (Figure 2A **& 3**), concurrently with moderate MD and RD increment indicating enhanced water diffusivity along all axes of a fitted tensor. As for the case of 100 Gy, the coincidence of RD increment and AD decrement at 180 d suggests a serious radiation-caused injury by exaggerating deleterious processes affecting white matter. The radiation necrosis further caused a local cell death, therefore a reduction in membrane restriction to water diffusion introduced a dramatic elevation to MD, RD as well as AD at 270 d, when a resolution of edema was observed (Figure 2A). An alternative but highly concurrent source for aberrant elevation in all DTI indices except for FA was the partial volume effect which biases signal toward the cerebrospinal fluid (CSF) in the enlarged ventricle due to cell loss. These effect hold still at 360 d though the AD came back to the level comparable to the baseline. Taken together, the alteration in these DTI indices indicates that SRS dose larger than or equal to 60 Gy would evoke persistent structural damages which took a half year to appear, whereas a small dose like 40 Gy or below caused no apparent harm in the surgical site.

**Figure 4.**
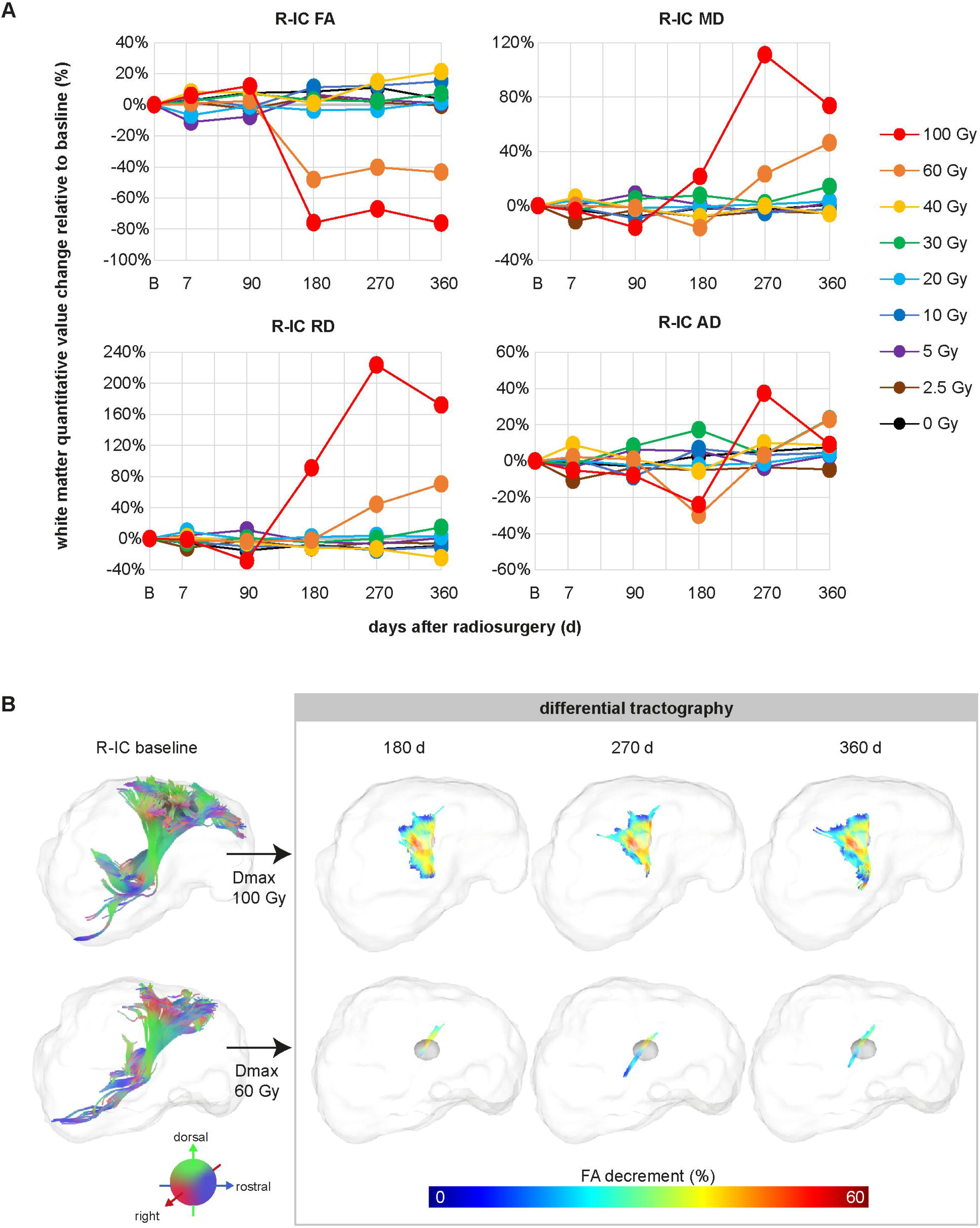
DTI indices analyses and injured tracts mapped by differential tractography. (A) The dose of 100 and 60 Gy caused white matter damages observed 180 days after surgery by showing a decreased FA, elevated MD and RD. The AD decrement reflecting axonal degradation was detected at 180 d in the dose of 100 Gy but at 270 d in 60 Gy, indicating a more deleterious process was induced by higher dose. The DTI indices were normalized to the pre-surgery baseline and expressed in percentage to enable comparison among different dose and across time. (B) The tract bundle passing through the planned surgical target in the baseline scan is illustrated by conventional tractography with colors denoting fiber orientation in the far left side. Differential tractography in the right box shows the SRS-injured tracts with color denoting the extent of FA decrement. The criterion for tracing difference between a specific time point and baseline was set at FA decrement > 30% and tract length > 10 mm. The results show that 100 Gy evoked diffused white matter injury while 60 Gy produced a local damage in few tracts. The position of R- IC ROI was noted by a small ball in the 3D rendering brain. FA for fractional anisotropy, MD for mean diffusivity, RD for radial diffusivity, AD for axial diffusivity.

The extent of radiation-caused axonal injury at a time was mapped by differential tractography which reveals tracts with segments showing FA decrement in comparison with the pre-surgery baseline (see **Figure S2** for example). The tract bundle of the right internal capsule passing through the planned surgery target is shown in the far left side of Figure 4B by conventional tractography, which reconstructs fiber tracts according to the principle diffusion orientation with green denoting dorso-ventral, red for right-left, and blue for rostro-caudal. The right box of Figure 4B illustrates the tracts showing FA decrement mapped at 180, 270, as well as 360 d after surgery with a color spectrum denoting FA change in percentage. These results show that the dose of 100 Gy evoked a profound white matter injury extending widely from the surgery site, while 60 Gy produced local impairment in few tracts passing through the surgical site. It is noticeable that distance-dependent FA reduction was observed relative to the surgical site with stronger toward proximal and mild toward distal. **Table 1** summarizes the SRS dose implemented in each pig and the number of tracts-with-difference in each time point. A long-term white matter damage without recovery from 180 d to 360 d was observed in the dose of 60 and 100 Gy as the number of tract-with-difference remained the same level. No tracts with obvious FA decrement were found in the dose lower than 60 Gy and control animals, while the result of 30 Gy at 360 d (tract number = 8) was ignored as a false positive result was found at 7 d (tract number = 57) when a structural change was not supposed to occur.

**Table 1.** SRS dose and the number of tracts with FA decrement through R-IC.

| Pigs | SRS dose |  | tracts with difference<sup>a</sup> in follow-up time points |  |  |  |  |
| --- | --- | --- | --- | --- | --- | --- | --- |
|  | L-M1 | R-IC | 7 d | 90 d | 270 d | 270 d | 360 d |
| #1 | 120 Gy | 100 Gy | 0 | 0 | 1921 | 1511 | 2015 |
| #2 | 100 Gy | 60 Gy | 0 | 0 | 28 | 34 | 26 |
| #3 | 80 Gy | 40 Gy | 0 | 0 | 0 | n.a.<sup>b</sup> | 0 |
| #4 | 60 Gy | 30 Gy | 57 | 0 | 0 | 0 | 8 |
| #5 | 40 Gy | 20 Gy | 2 | 0 | 0 | 0 | 0 |
| #6 | 30 Gy | 10 Gy | 0 | 0 | 0 | 0 | 0 |
| #7 | 20 Gy | 5 Gy | 0 | 0 | 0 | 0 | 0 |
| #8 | 10 Gy | 2.5 Gy | 0 | 0 | 0 | 0 | n.a.<sup>b</sup> |
| #9-11<sup>c</sup> | 0 Gy | 0 Gy | 0 | 0 | 0 | 0 | 0 |

<sup>a</sup>. the criterion for differential tractography was set at FA decrement > 30 %, fiber length > 10 mm
<sup>b</sup>. n.a. for analysis not available due to failure in image alignment toward the baseline scan
<sup>c</sup>. the results of three control animals were averaged

## Discussion

This research investigated the effect of stereotaxic radiosurgery on a white matter tract in the porcine brain to verify the radiation-induced structural and metabolic alteration. The right internal capsule was selected to receive focal irradiation, and anatomical MRI, DTI and FDG-PET scans were obtained once per three months for one year. The dose of 60 and 100 Gy produced radio-necrosis observed 180 d after surgery with signal elevation in T2W and FLAIR indicating edema as well as abnormal Gd accumulation denoting BBB breakage. Though these pathological change gradually recovered at 360 d, the dose of 100 Gy further caused brain structure deformation and ventricle enlargement. On the contrary, no apparent structural change was observed in the dose of 40 Gy or below. A small, non-necrotic dose may have a potential to be implemented in neuromodulation as a persistent local inhibition in brain metabolism was observed in some pigs, even a negative correlation between dose and SUV signal ratio was found at 360 d. The indices derived from revealed an obvious white matter injury induced by 60 and 100 Gy through reduction in FA and elevation in RD and MD. Via comparison in FA between time points, differential tractography further illustrates 100 Gy caused a diffused tract injury extending widely from the surgical site while 60 Gy only produced a local impairment within few tracts. In conclusion, a dose equal to or higher than 60 Gy applying onto the internal capsule was necrotic, and a period close to 180 d was needed to observe white matter injury. The dose of 40 Gy and below caused no apparent harm to brain structure, and future study was needed to verify the neuromodulatry effect.

Radiosurgery treatment for some diseases uses a necrotic dose on purpose to damage the specific tract that conveys signal in a hyper-active neural network. For example, obsessive compulsive disorder (OCD) was thought to have abnormal connectivity within the cortico-striato-thalamo-cortical circuit (Pauls et al., 2014), and the anterior limb of the IC was treated bilaterally to ameliorate symptoms. Trigeminal neuralgia, a chronic neuropathic facial pain, was commonly treated by applying focal radiosurgery at the trigeminal nerve root to disrupt transmission of pain signals to the brain (Hodaie et al., 2012; Regis et al., 2010; Tuleasca et al., 2019). The present study applied focal radiation onto the left M1 and right IC of normal healthy pigs, and white matter injury was detected in the right hemisphere when a dose of 60 Gy or above was selected. No specific behavioral task was applied on pigs in this study, but veterinary neurologic examination had been conducted frequently to monitor their sensorimotor function. Only the one receiving highest dose bilaterally (120 Gy for L-M1, 100 Gy for R- IC) showed aberration in gait and posture. Ataxia was first denoted in this pig from 180 d after surgery, and abnormal postural reaction like proprioception loss and deficit in single leg hopping was especially observed in the left side, suggesting the sensorimotor function is disturbed mainly in the right hemisphere. Amelioration of these malfunctions was found at 360 d in concurrence with the resolution of edema. In contrast, the pig receiving 60 Gy in R-IC appeared normal in movement all the time even it did have radiation necrosis and white matter injury detected a half year after surgery. These results indicate that the maximum dose of 60 Gy can provide a promising effect in down-regulating neural transmission by interfering white matter integrity locally but leaving subject’s sensorimotor ability undisturbed. However, a maximum dose of 160 Gy was usually selected for capsulotomy in treating OCD (Leveque et al., 2013; Miguel et al., 2019). The discrepancy in radiation dose used in literature and present study may come from the system adopted, as Gamma Knife was often reported yet CyberKnife was used in the present study. A seminal systemic review of trigeminal neuralgia treatment pointed out that the maximal doses in Gamma Knife SRS were 71.1-90.1 Gy prescribed at 100% isodose line and the ones in CyberKnife SRS were 64.3-80.5 Gy at 80% isodose line (Tuleasca et al., 2019). As the dose prescription was performed in a dissimilar fashion among radiosurgery techniques, deviation in maximum dose is inevitable when taking target selection and treatment time into account. A systemic review on clinical cases of capsulotomy is necessary in the future to provide recommendations in dose selection for different surgical systems to ensure safety and effectiveness. Furthermore, the study implementing porcine model for clinical translation can shed light on the dose selection if a behavioral paradigm bearing a resemblance to OCD is built.

The deleterious effect of radiation on white matter can be measured by DTI quantitative indices reflecting the extent of water diffusion. Either focal irradiation like what applied in the present study or radiotherapy prescribed to treat brain tumors caused white matter damage in a dose-dependent manner exhibiting FA decrement as well as RD and MD increment (Connor et al., 2017). Intact white matter fibers can be reconstructed through tractography techniques, while an FA value fall-off will lead to tract pruning which can only be inspected visually. For example, the tracts of trigeminal nerve were pruned in the patients responsive to radiosurgery (Tohyama et al., 2018) or at the site receiving focal radiosurgery (Hodaie et al., 2012). In the present study, a novel technique named differential tractography was adopted via repeat DTI scans to identify the exact segment of tracks illustrating FA reduction, by which a quantitative and objective measurement is provided to monitor the degree of injury. As shown in Figure 4B, an obvious dose- and distance-dependent effect was illustrated by a higher dose rendering profound injury and tract segments proximal to the surgical site exhibiting stronger FA reduction. There may be concerns about false positive results derived from FA comparison across time, but this can be mitigated through data quality control steps embedded in the analyzing software (DSI studio) and a conservative FDR determined by researchers. Since no reversed phase-encoding dataset was obtained in the present study to correct potential susceptibility-induced distortion, the data passing quality control steps were inspected visually with carefulness and two datasets were excluded from analysis due to unsuccessful image alignment. We have a high level of confidence in the current results, because we adopted a common criterion in detecting tract-with-difference among pigs. A stringent FDR lower than 0.01 were selected under the threshold examination in FA change and length on the pig (#1) showing the most prominent radiation induced change where tracts-with-difference detected at 7-d post-surgery were seen as false recovery (**Table S1**). Another concern about the result is that a standard b value of 1000 s/mm^2^ was used in present study, while a high b value like 4000 s/mm^2^ was reported to have higher sensitivity in detecting white matter difference (Baumann et al., 2012). In the meanwhile, high b-value acquisition was strongly recommended by Yeh et al. (2019), the inventors of differential tractography, to increase the power in detecting early neuronal injury and to prevent false-positive results. DTI at high b-values is sensitive to slow, restricted diffusion in the intracellular axonal space, whereas that at low b-values reflects fast diffusion owing to free water in the extracellular space. Regarding the deleterious effect of radiation revealed by DTI indices, Connor et al. (2016) utilized DTI with multiple b-values and found that greater radiation-induced changes were seen at lower b-values than those derived at higher b-value. A similar finding reported by Karunamuni et al. (2017) indicates that dose-dependent change was observed in the fast component of diffusion while dose-independent change was observed in the slow component. Based on the evidence above, DTI metrics derived from a b value like 1000 s/mm^2^ can reveal white matter injury in a dose-dependent manner as seen in our study with dose of 60 and 100 Gy. It is worth further exploration to examine whether DTI with higher b values can enhance sensitivity in identifying slow component diffusion in the normal appearing white matter treated with 2.5-40 Gy.

Long being perceived as a surgical technique that ablates tissues non-invasively, radiosurgery can also serve as a neuromodulation tool by applying sub-lethal radiation focally (Regis et al., 2010; Schneider et al., 2010; Schneider et al., 2021). Multiple lines of evidence showed that low dose irradiation can regulate brain activity without tissue destruction in porcine model. A small dose like 40 Gy caused no apparent harm in gray matter structures observed from MRI (Zaer et al., 2020), and neither vascular change nor radiation-induced inflammation reaction was found in the histological examination (Zaer et al., 2022). An increased 18F-FDG-uptake was detected in the L-M1 treated with 10-40 Gy, and this effect appeared at 180 d and became even evident at 270 d after surgery (Yeh et al., 2021). Electrophysiological recording further illustrated an increased spontaneous firing rate and a decreased visual evoked potential (VEP) P1 peak time in the 40 Gy-treated visual cortex (Zaer et al., 2022), with which the former indicates higher basal activity and the later reflects faster information processing in the visual circuit. The neuromodulatory effect may be achieved by an increment in neuronal excitability, and this local change further tunes the activity balance within, but not limited to, a certain functional circuit to cause a long-term modulation. As a result, an enduring alteration in the brain activity marker, like 18F-FDG uptake, would thus be detected. Nevertheless, the mechanism of how low dose irradiation increases the neuronal activity and why this effect requires at least a half year to develop remain to be elucidated in the future.

Notably, in the present study a non-necrotic low dose had a potential to inhibit, but not to enhance, 18F-FDG uptake in the surgical target located in the R-IC. The reversal effect in neuromodulation can be explained by the cell constitution in the target area, as the white matter contains a higher glia-to-neuron ratio than gray matter. Glia and their progenitors were more susceptible to radiation than neurons (Belka et al., 2001; Hua et al., 2012), such that irradiation may impede local metabolism by interfering gliogenesis or proliferation especially in the white matter areas. Lower dose may promote neuronal excitability and cause a plausible decrement in metabolic activity of glia simultaneously, and the amount of 18F-FDG uptake would thus reflect the summation of these two independent processes. As a result, the same dose would have opposite effects when being applied onto a cortical area and a tract, respectively, with activity alteration contributed from neuronal activation dominates that from functional suppression of glia or vice versa. Under this scenario, the local metabolism in the R-IC is expected to negatively correlate with the dose as the lower the radiation the less the interference to glial function. The result from 360 d after surgery did confirm this speculation. In comparison to the abundant signal level in the gray matter, the lower 18F-FDG absorption rate in the white matter zone makes it less sensitive to detecting subtle changes. An alternative approach for assessing the modulatory effect is to examine the regions connected by the R-IC but not the tract per se. With the attempt to explore every potential region where neuromodulation might occur, conventional tractography was adopted by planting seeds in the surgical area to map the regions having afferents/efferents passing through. Regions like the primary somatosensory cortex, M1, thalamus, and pons were examined, but to our surprise no clear effect was found to draw a conclusion about the dose-dependent response (data not shown). A plausible explanation for this negative finding is that the modulatory effect on a white matter tract exerts on signal interplay between regions rather than on local metabolic activity, thus a static PET scan measuring the distribution of radiotracer at a particular moment is not able to capture this dynamic interaction. Resting state fMRI is a good tool to detect the functional connectivity (FC) change by calculating the signal fluctuation between regions and reveal the interaction among functional networks (Biswal et al., 1995), when suitable agents for animal anesthetization were adopted (Simchick et al., 2021; Sirmpilatze et al., 2019). If a FC change bear a correlation with a behavioral alteration after SRS, the network nodes are the key substrates where neuromodulation occur.

This study utilized a miniature pig model which has a gyri pattern as complex as human brain to investigate the effect of stereotaxic surgery applied to white matter by a longitudinal study with anatomical MRI, DTI, as well as FDG-PET. Administration of doses equal to or higher than 60 Gy led to a late-onset radionecrosis which took a period close to 180 d to develop edema and BBB breakage. The extent of white matter injury was identified through differential tractography denoting FA reduction in comparison to pre-surgery baseline. The result indicates that 60 Gy produced a local impairment within few tracts, by which the information transmission in a hyper-active network can thus be down-regulated. In contrast, doses of 40 Gy and below did not result in any discernible harm to the brain structure, and a sustained local inhibition in brain metabolism was observed in some pigs. When employing radiosurgery in capsulotomy, the maximum dose needs to be considered carefully, given the way in dose prescription was dissimilar across radiosurgery techniques. Regarding the potential application in neuromodulation, a comprehensive assessment throughout the whole brain is necessary to evaluate its effect. The derived brain index must be correlated with changes in behavior or cognitive function to confirm the neuro-modulatory effect and to identify the critical neural substrate responsible for its occurrence.

## Conclusion

Stereotaxic radiosurgery delivered focally onto the internal capsule in a pig brain can induce white matter injury when a dose of 60 Gy or above was employed. The effect of neuromodulation by low-dose radiation awaits a comprehensive assessment throughout the whole brain in the future.

## Acknowledge

The authors would like to thank the Taiwan Instrument Research Institute (TIRI) and National Taiwan University Hospital for the support in MRI and FDG-PET experiments respectively.

## Supplementary materials

Table S1 illustrates the threshold test for FA change (%) and tract length (mm) derived from the pig #2, as it showed prominent radiation-induced change in the anatomical MRI at 360 d after surgery. White matter structure change took time to occur, therefore what found in the dataset of 7d was seen as false discovery. Number of tracts-with-difference was denoted under various threshold for differential tractography, and the criterion of FA reduction larger than 30% with tract length longer than 10 mm was selected to control the false discovery rate (FDR) lower than 0.01.

Figure S1 shows the original value of derived DTI indices, and the indices of L-IC serving as the negative control illustrate no change across time. Figure S2 provides an example of the image aligned for differential tractography. The result of image alignment and the location of the surgical target were visually inspected with carefulness.

*[insert Fig S1 here]*

Figure S1. The DTI matrices for right and left internal capsule.

*[insert Fig S2 here]*

Figure S2. Example of image alignment of follow-up images toward pre-surgery baseline. The white square illustrates the surgical target in the right internal capsule. The stick represent the reconstructed fiber and color denotes orientation.

---

**Table S1.** Threshold test for FA reduction (%) & tract length (mm) and the resulting FDR.

| Tracts showing FA decrement in 7 d compared to pre-surgery baseline |  |  |  |  |  |
| --- | --- | --- | --- | --- | --- |
|  | 5 mm | 10 mm | 15 mm | 20 mm | 25 mm |
| 10% | 43851 | 10869 | 3726 | 1252 | 422 |
| 15% | 19610 | 4296 | 477 | 0 | 0 |
| 20% | 8710 | 1131 | 13 | 0 | 0 |
| 25% | 3546 | 215 | 0 | 0 | 0 |
| 30% | 1680 | 40 | 0 | 0 | 0 |
| 35% | 928 | 21 | 0 | 0 | 0 |
| 40% | 453 | 0 | 0 | 0 | 0 |

| Tracts showing FA decrement in 360 d compared to pre-surgery baseline |  |  |  |  |  |
| --- | --- | --- | --- | --- | --- |
|  | 5 mm | 10 mm | 15 mm | 20 mm | 25 mm |
| 10% | 92016 | 39766 | 19460 | 9591 | 4106 |
| 15% | 53499 | 23021 | 10815 | 4981 | 1475 |
| 20% | 31366 | 12899 | 5405 | 2145 | 398 |
| 25% | 19054 | 7643 | 3149 | 774 | 97 |
| 30% | 11782 | 4857 | 1621 | 385 | 0 |
| 35% | 8363 | 3347 | 1024 | 116 | 0 |
| 40% | 5366 | 1893 | 383 | 0 | 0 |

| False discovery rate (FDR) = # of tracts found in 7 d over that in 360 d |  |  |  |  |  |
| --- | --- | --- | --- | --- | --- |
|  | 5 mm | 10 mm | 15 mm | 20 mm | 25 mm |
| 10% | 0.477 | 0.273 | 0.191 | 0.131 | 0.103 |
| 15% | 0.367 | 0.187 | 0.044 | 0.000 | 0.000 |
| 20% | 0.278 | 0.088 | 0.002 | 0.000 | 0.000 |
| 25% | 0.186 | 0.028 | 0.000 | 0.000 | 0.000 |
| 30% | 0.143 | **0.008** | 0.000 | 0.000 |  |
| 35% | 0.111 | 0.006 | 0.000 | 0.000 |  |
| 40% | 0.084 | 0.000 | 0.000 |  |  |

