## Supplementary figures and images for "Tracking White Matter Changes After Stereotaxic Radiosurgery in Miniature Pigs Using structural MRI, DTI, and FDG-PET"

### Figure S1

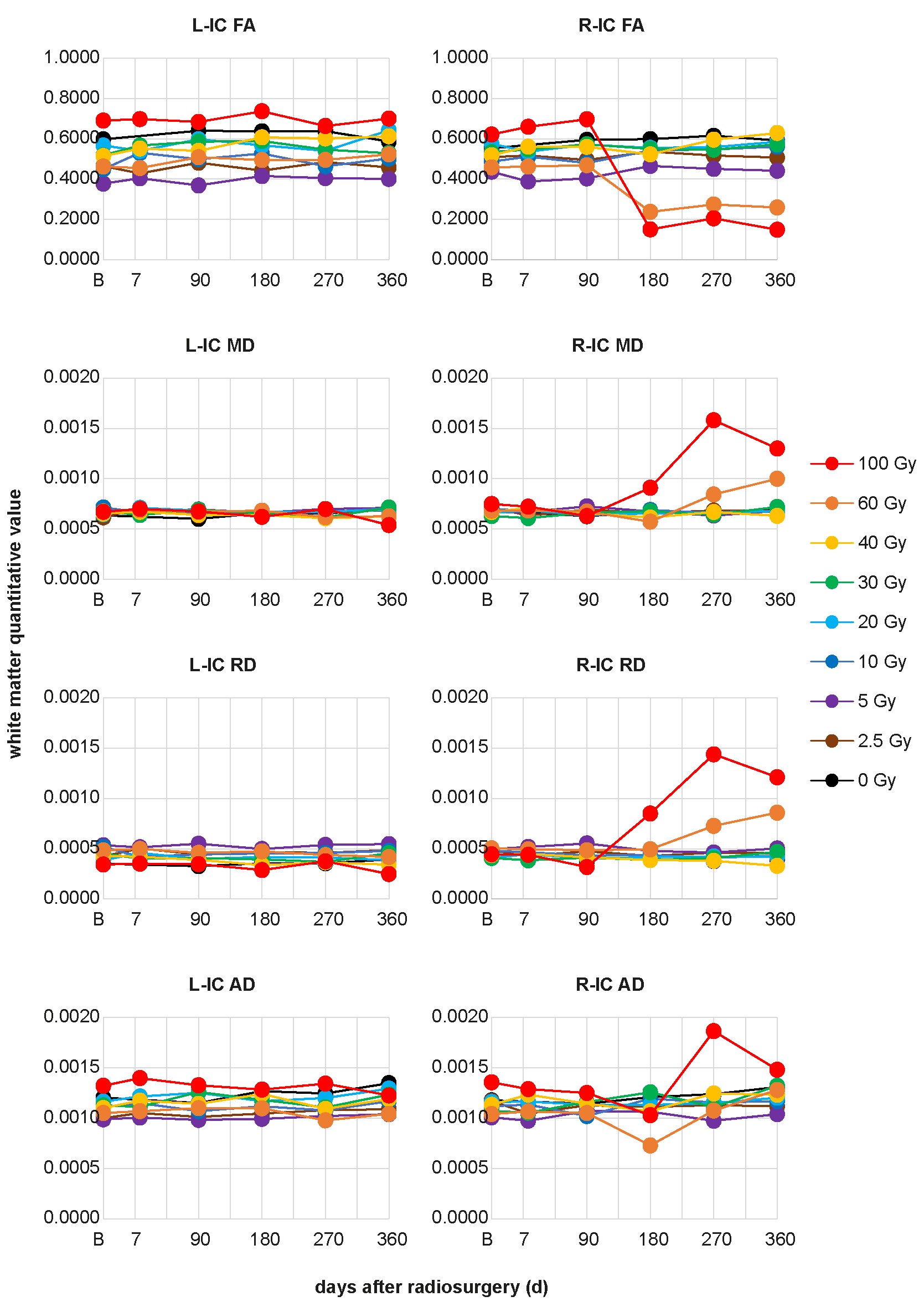

### Figure S2

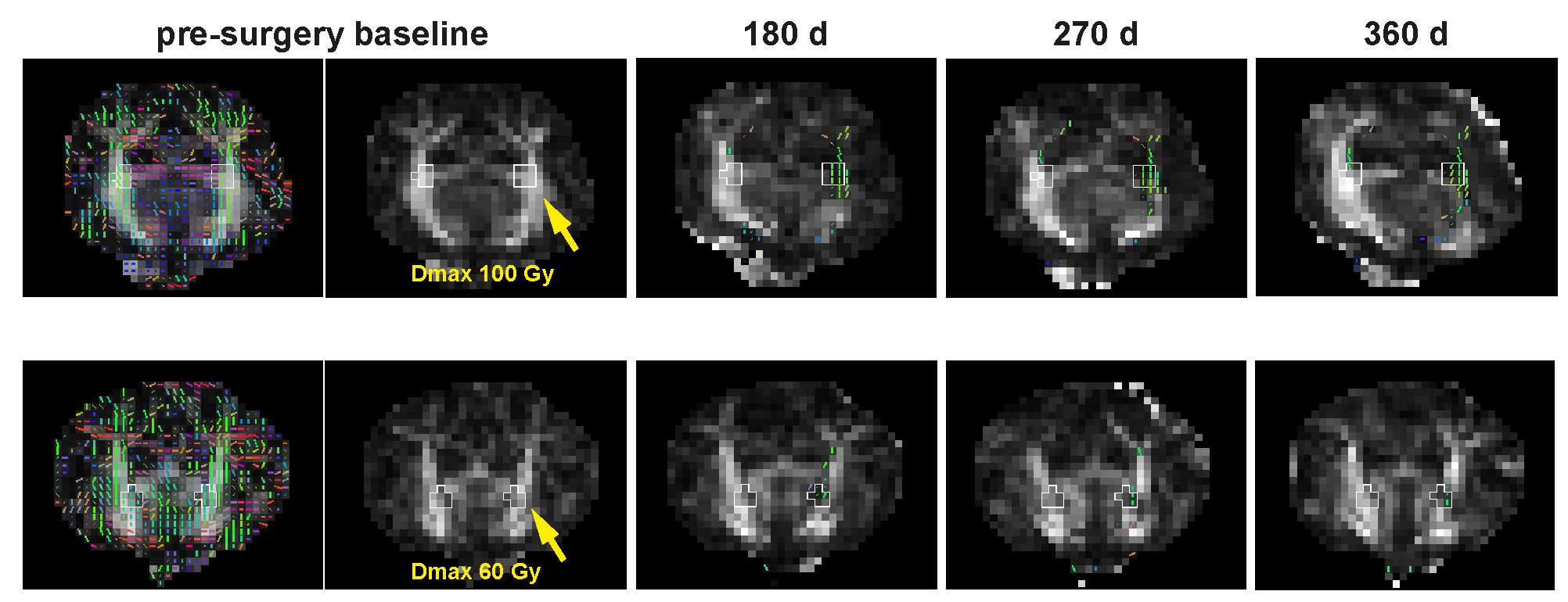
